# FAXC and Metaxin Proteins of Fungi and Bacteria: Distinct Categories of Proteins with Shared Structural Features

**DOI:** 10.1101/2025.01.06.631459

**Authors:** Kenneth W. Adolph

## Abstract

FAXC (failed axon connections) proteins are shown to be present in fungi and bacteria and to have structural homology to metaxin proteins of vertebrates and invertebrates. Fungal and bacterial FAXC proteins also share structural features with the metaxin-like proteins of plants and protists. But the FAXC and metaxin proteins are different categories of proteins, as seen in the low percentages of identical amino acids in pairwise sequence alignments. The metaxin-like structural features include the conserved protein domains that are characteristic of metaxin proteins. These are the GST_N_Metaxin, GST_C_Metaxin, and Tom37 domains. Another metaxin-like structural feature is a conserved pattern of α-helical and β-strand secondary structure. The FAXC proteins of both fungi and bacteria share this pattern of secondary structure with vertebrate and invertebrate metaxin proteins. The predicted 3D structures of fungal and bacterial FAXC proteins revealed a characteristic 4-stranded, β-sheet motif that was also found to characterize metaxin proteins. Phylogenetic analysis demonstrated that fungal and bacterial FAXC proteins are related by evolution to vertebrate metaxin proteins, though more closely related to vertebrate FAXC proteins. The neighboring genes of the FAXC genes of fungi are different than the neighboring genes of vertebrate FAXC and metaxin genes. A high level of protein sequence homology was found for the multiple predicted FAXC proteins of fungi that have multiple FAXC genes, and in comparing the FAXC proteins of different species of bacteria of the same genus.

## 1. INTRODUCTION

The *fax* (failed axon connections) gene was first described in *Drosophila melanogaster* as a novel gene that is primarily expressed in nerve axons (Hill et al., 1995). Further studies of the role of the Fax protein in *Drosophila* or other species have not been reported in the literature. However, genes homologous to the *Drosophila* gene have been found to be widely distributed in a variety of vertebrates and invertebrates (Adolph, 2023). The predicted FAXC proteins of vertebrates and invertebrates have structural features that are characteristic of metaxin-like proteins. These features include metaxin-like protein domains, similar patterns of α-helical and β-strand secondary structure, and similar phylogenetic relationships. To expand upon these prior results (Adolph, 2023), the focus of this study was to further investigate the metaxin-like structural features of the FAXC proteins of fungi and bacteria.

The human FAXC homolog of the *Drosophila fax* gene, also known as the *C6Orf168* gene (Chromosome 6 Open Reading Frame 168), is at cytogenetic location 6q16.2. Vertebrate FAXC genes, in addition to human, that are used in this report for comparison with fungal and bacterial FAXC genes are those of the mouse, zebrafish, and frog. These vertebrates were chosen because they are widely studied model systems and have fully sequenced genomes (mouse: Mouse Genome Sequencing Consortium, 2002; zebrafish: Howe et al., 2013; frog: Session et al., 2016). Numerous invertebrates of various phyla also have genes homologous to the *Drosophila fax* gene. The invertebrate phyla include Arthropoda, Mollusca, Cnidaria, and Placozoa. Examples of invertebrates with FAXC genes are the Atlantic horseshoe crab *Limulus polyphemus*, the California sea hare *Aplysia californica*, the starlet sea anemone *Nematostella vectensis*, and the placozoan *Trichoplax adhaerens*.

A possible role for FAXC proteins of vertebrates and invertebrates is in axonal development, in view of the results with *Drosophila*. However, the similar structural features of FAXC and metaxin proteins suggest that the role of FAXC proteins should be considered in relation to the role of the metaxins. The similar structural features include conserved protein domains, secondary structures, and 3D structures predicted with AlphaFold (Jumper et al., 2021). Experimental evidence originally revealed that metaxin 1 of mouse and human is a protein of the outer mitochondrial membrane with a role in the uptake of new proteins into mitochondria. Metaxin 2 was subsequently identified as a vertebrate protein working along with metaxin 1 in protein import into mitochondria. Metaxin 3 was more recently described and shown to have a wide occurrence among vertebrates, though its role is unknown.

Metaxin 1 and 2 homologs have also been identified in invertebrates, including the model research organisms *Drosophila melanogaster* and *C*. *elegans* (Adolph, 2020a). Metaxins are not confined to vertebrates and invertebrates, however, and homologous proteins have been found in plants such as the model plant of molecular biology research *Arabidopsis thaliana* and the economically important plant *Glycine max*, the soybean (Adolph 2020b). In addition, metaxin-like proteins are present in protists, which are eukaryotes that are not animals, plants, or fungi, but a separate category (Adolph, 2021). Among the protists are the human malaria parasite *Plasmodium vivax* and the plant pathogen *Phytophthora infestans*.

Fungi with FAXC genes and completely sequenced genomes were used for this study. These include three fungi of the Neocallimastigomycota division: *Neocallimastix californiae*, *Piromyces finnis*, and *Anaeromyces robustus* (genome sequences: Haitjema et al., 2017; Mondo et al., 2017). They are anaerobic symbionts of large herbivores. The FAXCs of three fungi of the Chytridiomycota division were also part of this investigation: *Gonapodya prolifera* (Chang et al., 2015), *Powellomyces hirtus* (Amses et al., 2022), and *Spizellomyces punctatus* (Russ et al., 2016). Another fungus selected for this study was *Heliocybe sulcata*, a mushroom of the Basidiomycota division (Varga et al., 2019).

The bacteria in this study that have FAXC genes and completely sequenced genomes include examples of a variety of bacterial phyla and classes. *Pseudomonas aeruginosa* (genome sequence: Stover et al., 2000) is one of the major causes of opportunistic infections in humans. It is in the Pseudomonadota phylum, Gammaproteobacteria class. *Rhodococcus* strains (phylum: Actinomycetota) are important because of their ability to degrade a wide range of organic molecules (McLeod et al., 2006). Another example of a bacterium with FAXC genes is *Rhodospirillum rubrum* (Munk et al., 2011), one of the most intensively investigated microbial species. It is in the Alphaproteobacteria class.

The purpose of this study was to examine the presence of FAXC genes in fungi and bacteria, and to determine fundamental properties of the predicted FAXC proteins. This follows the previous investigation of FAXC genes and proteins in vertebrates and invertebrates (Adolph, 2023). In the present study, homologs of FAXC genes were found to be widely distributed among both fungi and bacteria. A focus was upon the relationship between FAXC and metaxin proteins of fungi and bacteria, since FAXC proteins of vertebrates and invertebrates were found to be characterized by metaxin-like structural features (Adolph, 2023). The results of the present study show that FAXC proteins of fungi and bacteria also have metaxin-like structural features. The metaxin-like features of fungal and bacterial FAXC proteins include: (1) GST_N_ and GST_C_Metaxin protein domains, (2) conserved patterns of α-helical segments, and (3) a conserved 3D β-sheet motif. Further investigations should help to reveal additional aspects of both the structure and function of the FAXC proteins of fungi and bacteria.

## 2. METHODS

Pairs of protein sequences were aligned using the NCBI BLAST Global Align and Align Two Sequences tools (Figures 1 and 7) to determine the percentages of identical and similar amino acids (https://blast.ncbi.nlm.nih.gov/Blast.cgi; Needleman and Wunsch, 1970; Altschul et al., 1990). The NCBI CD search tool was employed to identify the conserved protein domains of FAXC proteins (www.ncbi.nlm.nih.gov/Structure/cdd/wrpsb.cgi; Lu et al., 2020). The major domains (GST_N_Metaxin, GST_C_Metaxin, and Tom37) of representative fungal and bacterial FAXC proteins are in Figure 2A and 2B. Vertebrate domains, including the FAXC_N domain, are in Figure 2C. The PSIPRED server (bioinf.cs.ucl.ac.uk/psipred/; Jones, 1999; Buchan and Jones, 2019) was utilized to predict the α-helical and β-strand segments of FAXC and metaxin proteins. Multiple-sequence alignments to compare the secondary structure segments were carried out with the NCBI COBALT alignment tool (www.ncbi.nlm.nih.gov/tools/cobalt;

**Figure 1.**
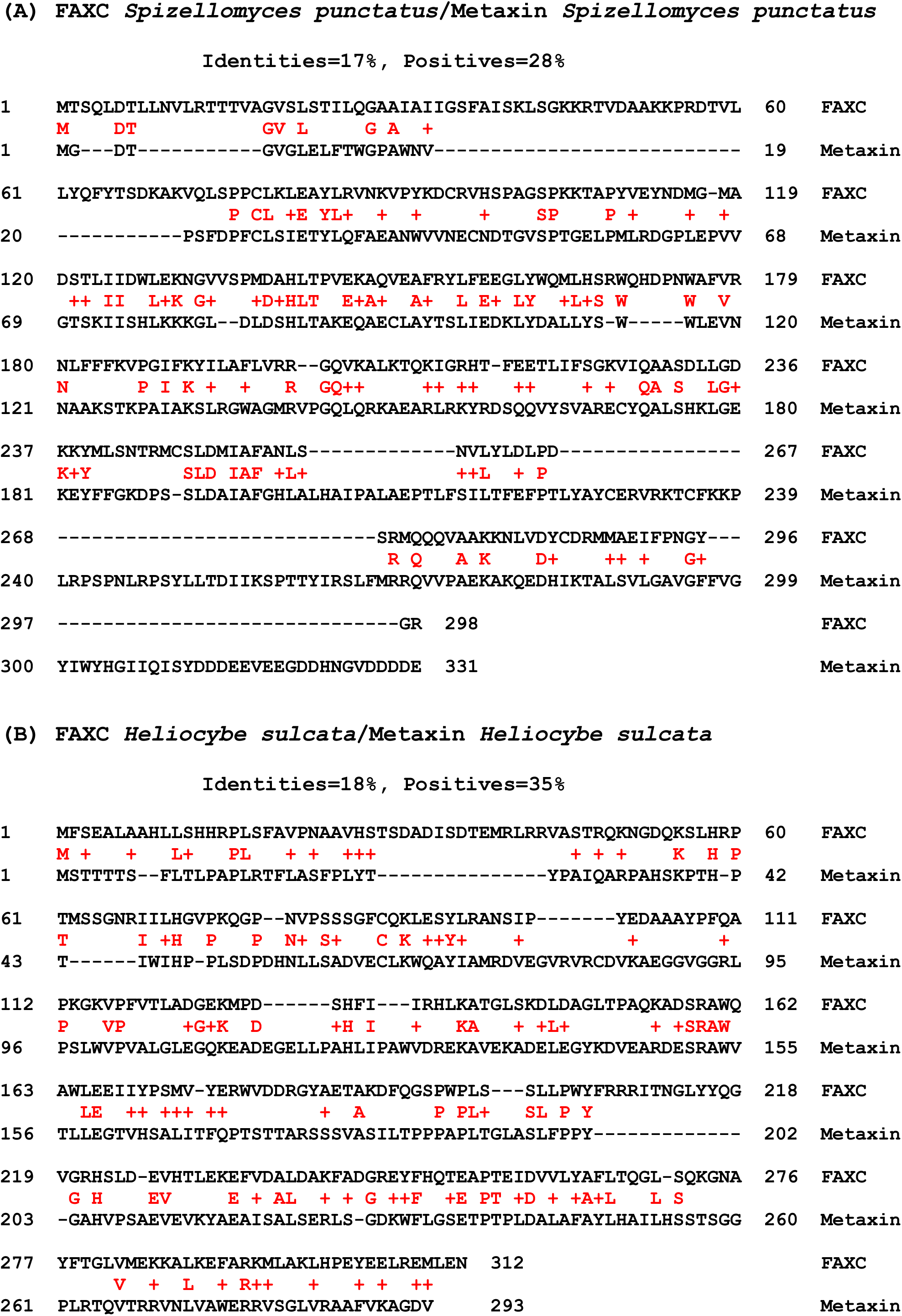
Protein sequence alignments of FAXC and metaxin proteins of representative fungi. In (A), the alignment includes the FAXC and metaxin amino acid sequences of the soil fungus *Spizellomyces punctatus* (taxonomic division: Chytridiomycota). The identical amino acids are 17% and the positives are 28%. The alignment used NCBI Global Align to compare the entire sequences. The low percentages demonstrate that the FAXC and metaxin proteins of this fungus are different categories of proteins. (B) shows the alignment of the FAXC and metaxin proteins of the basidiomycete mushroom *Heliocybe sulcata*. The identities are 18% and the positives, 35%. The low percentages again demonstrate that the FAXC and metaxin proteins are separate categories of proteins. Results like these were also found for a variety of other fungi.

**Figure 2.**
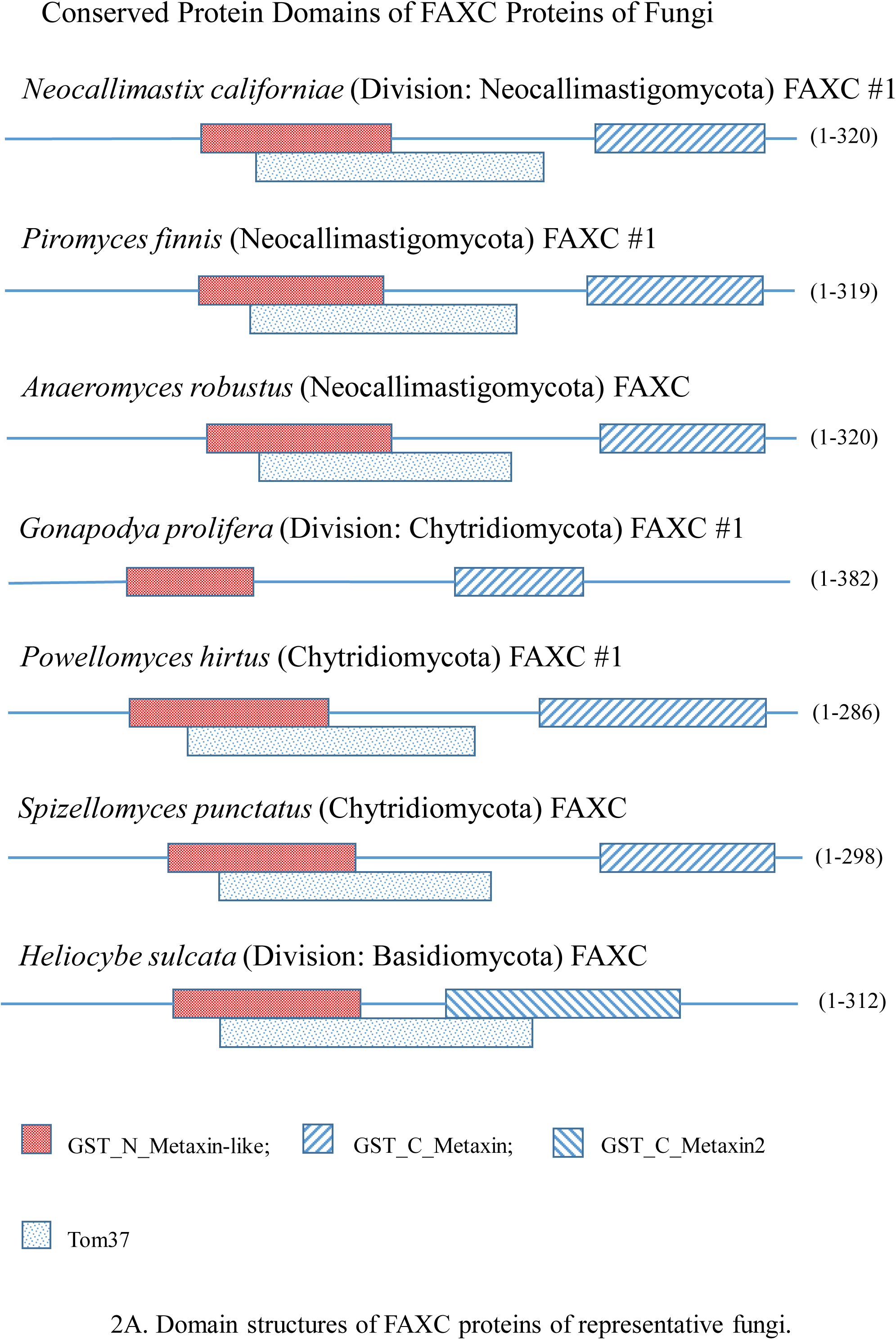
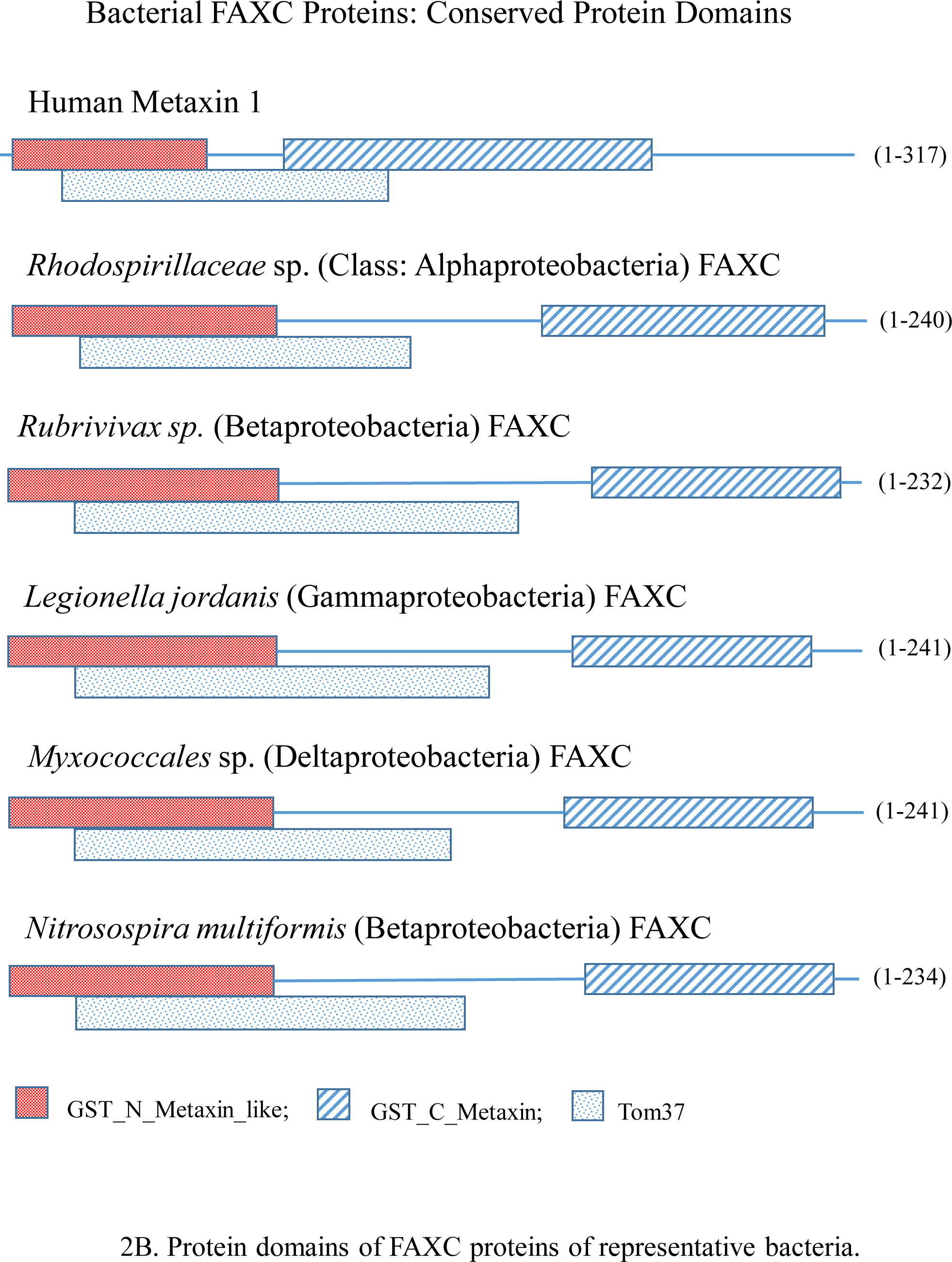
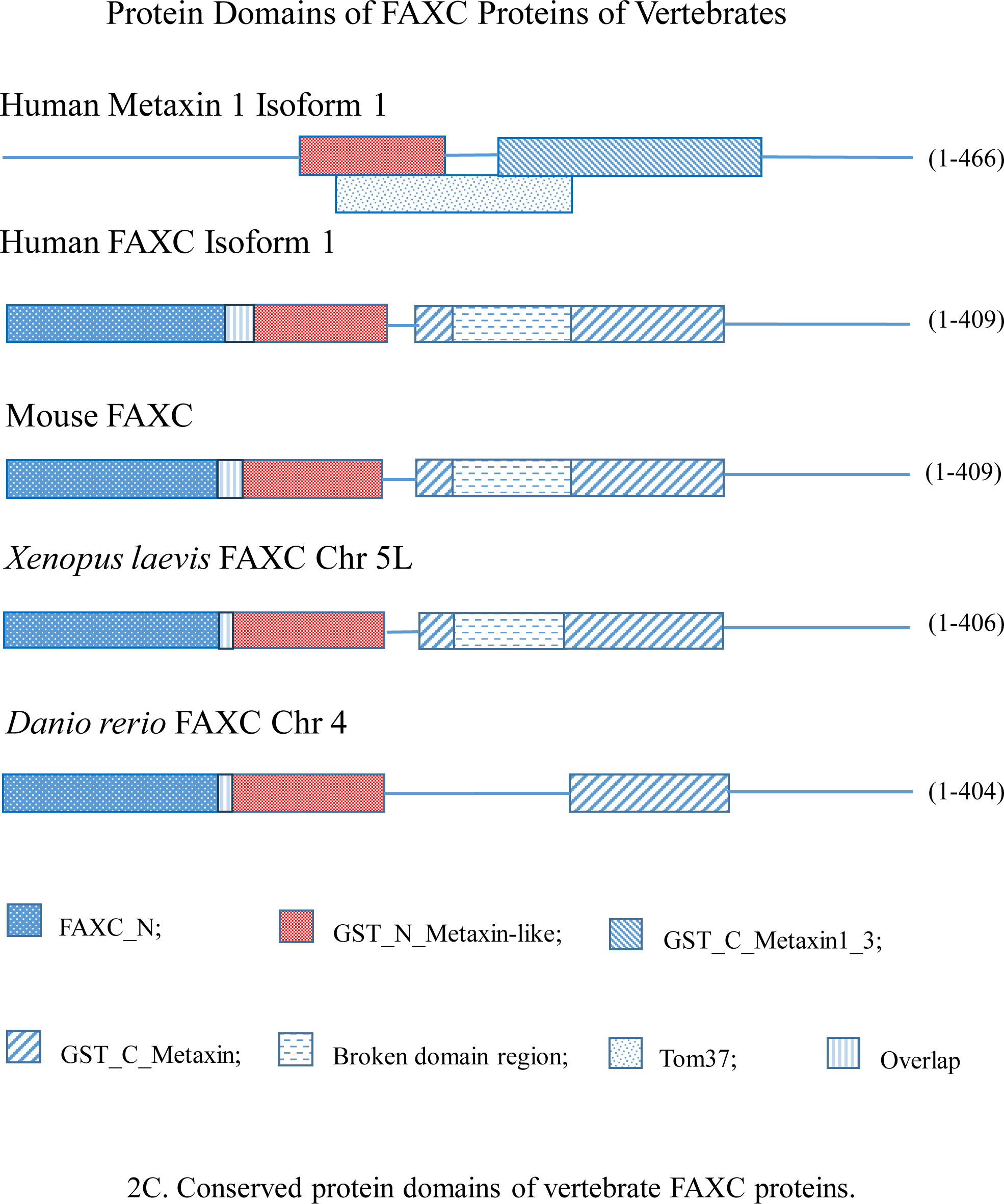
Conserved protein domains of fungal and bacterial FAXC proteins. (A) The GST_N_Metaxin and GST_C_Metaxin domains are shown for FAXC proteins of fungal divisions that include Neocallimastigomycota, Chytridiomycota, and Basidiomycota. The GST_N_Metaxin and GST_C_Metaxin domains are the major, characteristic protein domains first identified for the metaxin proteins of the mouse, humans, and other vertebrates. Also shown is the Tom37 domain, which is a common domain of metaxin proteins and metaxin-like proteins besides FAXC proteins. Fungi can have more than a single FAXC gene and predicted protein, and FAXC #1 protein is included in the figure for species with multiple predicted FAXC proteins. (B) The figure includes the major domains (GST_N_Metaxin, GST_C_Metaxin, and Tom37) that are characteristic of the FAXC proteins of bacteria. These are also the major domains of vertebrate metaxin proteins and FAXC proteins, such as human metaxin 1 in the figure. The bacteria shown are representative of four classes of the Pseudomonadota (Proteobacteria) phylum, in particular the Alphaproteobacteria, Betaproteobacteria, Deltaproteobacteria, and Gammaproteobacteria. (C) Protein domain structures of vertebrate FAXC proteins. In addition to the GST_N_Metaxin and GST_C_Metaxin domains, vertebrate FAXC proteins also possess the FAXC_N domain. The FAXC_N domain has only been identified in vertebrate FAXC proteins and is not present in fungi (Figure 2A), bacteria (Figure 2B), or in other species including plants and protists. The FAXC_N domain extends from the N-terminus of the FAXC proteins to the GST_N_Metaxin domain. There is a short overlap between the FAXC_N domain and the GST_N_Metaxin domain.

Papadopoulos and Agarwala, 2007). Examples are in Figure 3. Predicted 3D protein structures were obtained from the AlphaFold protein structure database (Jumper et al., 2021; Varadi et al., 2022). The structures (Figure 4) revealed a characteristic 4-stranded, 3D β-sheet motif for FAXC and metaxin proteins. The 3D structures were viewed with the iCn3D structure viewer associated with NCBI BLAST protein searches (Wang et al., 2020). Evolutionary relationships were examined with the COBALT multiple-sequence alignment tool and the phylogenetic trees generated using the alignments. Typical phylogenetic results are shown in Figure 5A and 5B for FAXC proteins of fungi and bacteria. Neighboring genes of FAXC and metaxin genes (Figure 6) were identified via the NCBI “Gene” database, and specifically through “Genomic regions, transcripts, and products”.

**Figure 3.**
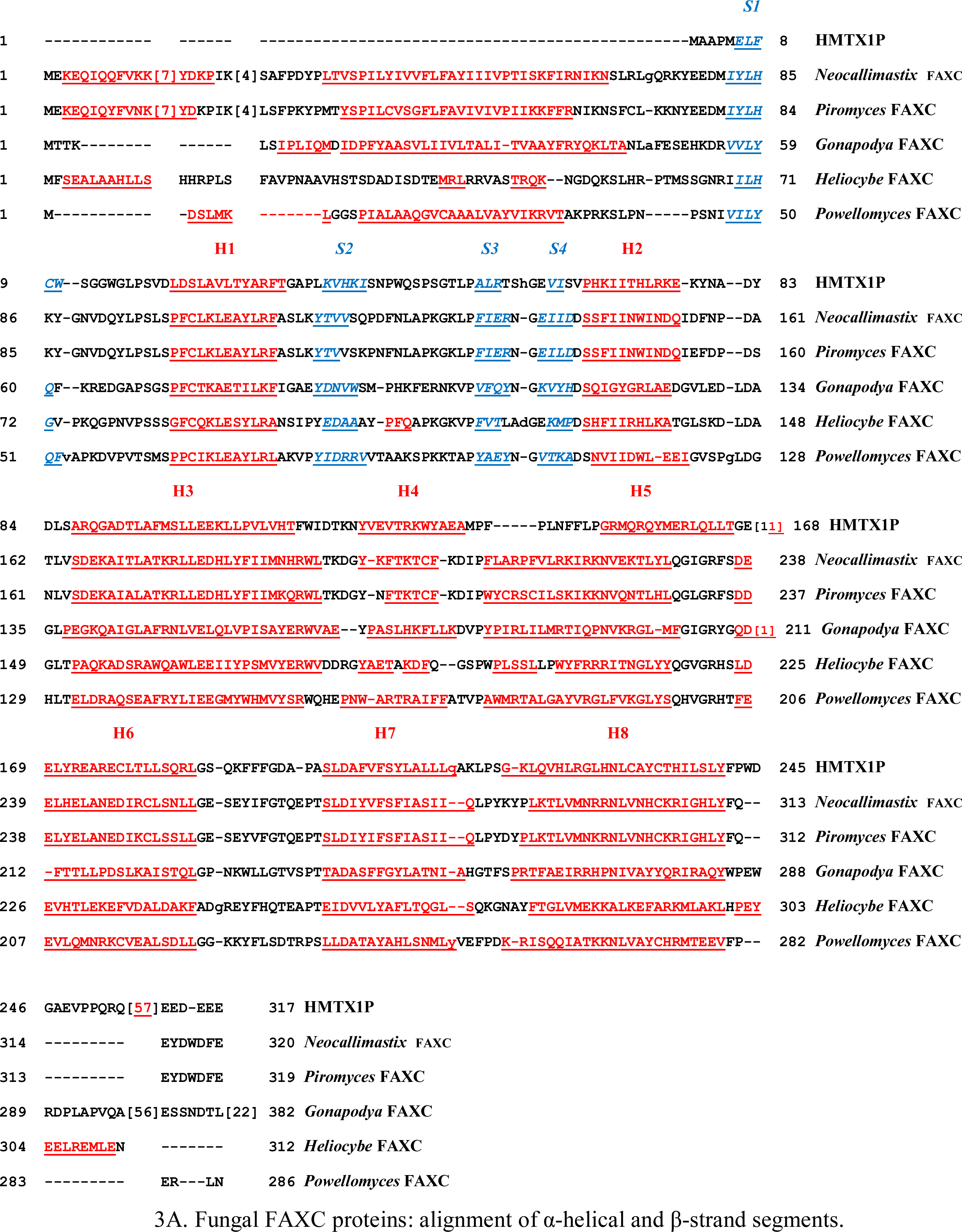
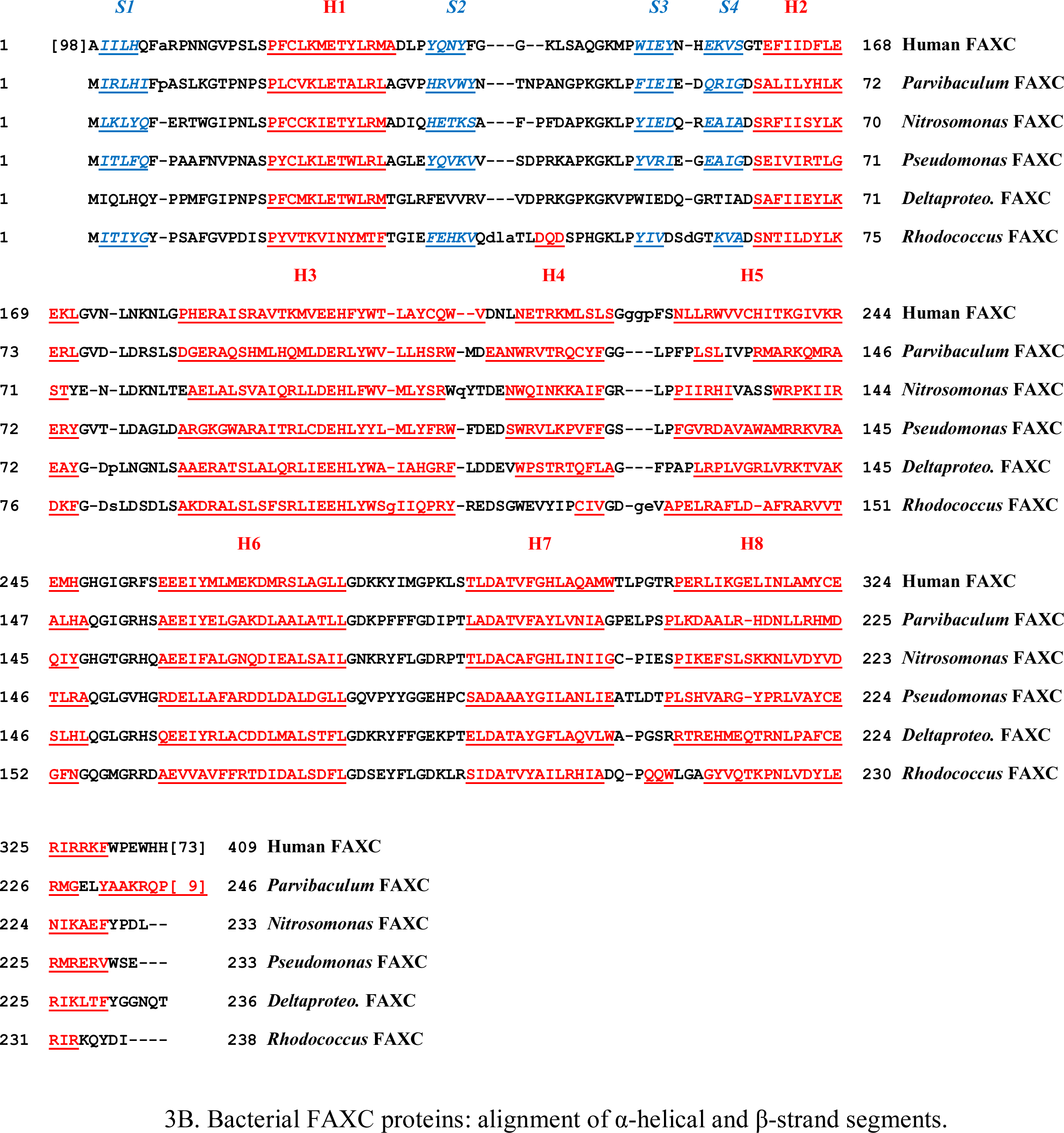
Secondary structure of FAXC proteins of fungi and bacteria: α-helical and β-strand segments. (A) Major features that dominate the predicted secondary structure of fungal FAXC proteins are a conserved pattern of α-helices and a conserved β-strand motif. The pattern of α-helices is almost identical in comparing the fungal FAXC proteins with human metaxin 1 (HMTX1P), and the same is true of the pattern of β-strand segments. In the figure, the α-helical segments have a red color and are underlined, while the β-strand segments are blue and are underlined and italicized. The α-helices are labeled H1 through H8 and the β-strand segments S1 through S4, following the labeling for human metaxin 1 protein. The fungal FAXC proteins have additional helices near their N-terminal ends, while human metaxin 1 has an extra helix H9 within [57] near its C-terminus. The β-strand segments S2 – S4 are between helices H1 and H2, and S1 is before H1. All of the fungi shown have two FAXC proteins, FAXC #1 and FAXC #2. The FAXC in the figure is FAXC #1. The fungi represent divisions that include Neocallimastigomycota (*Neocallimastix*, *Piromyces*), Chytridiomycota (*Gonapodya*, *Powellomyces*), and Basidiomycota (*Heliocybe*). (B) The predicted α-helical segments of FAXC proteins of representative bacteria are in red and underlined, and are labeled H1 through H8. The β-strand segments are blue, underlined, italicized, and labeled S1 through S4. The secondary structure is dominated by the helical segments, which are highly conserved among all of the bacterial FAXC proteins and human FAXC isoform 1. β-Strand segments are also highly conserved, with S2 – S4 between H1 and H2, and S1 at the N-termini of the bacterial proteins. The bacteria in the figure belong to the Pseudomonadota (Proteobacteria) phylum and the taxonomic classes Alphaproteobacteria (*Parvibaculum lavamentivorans*), Betaproteobacteria (*Nitrosomomas nitrosa*), Gammaproteobacteria (*Pseudomonas aeruginosa*), and Deltaproteobacteria. The Actinomycetota phylum is represented by a *Rhodococcus* sp. bacterium.

**Figure 4.**
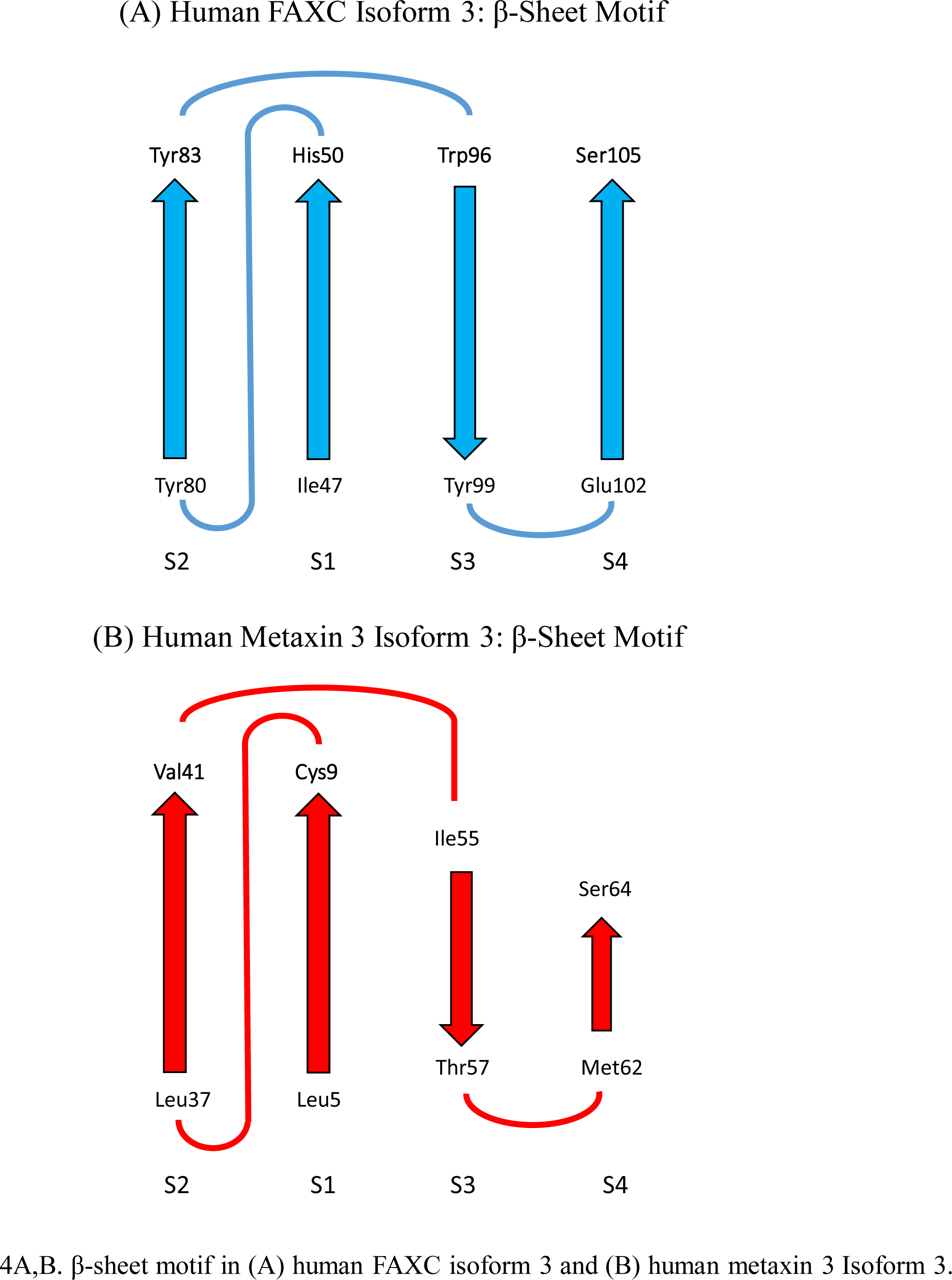
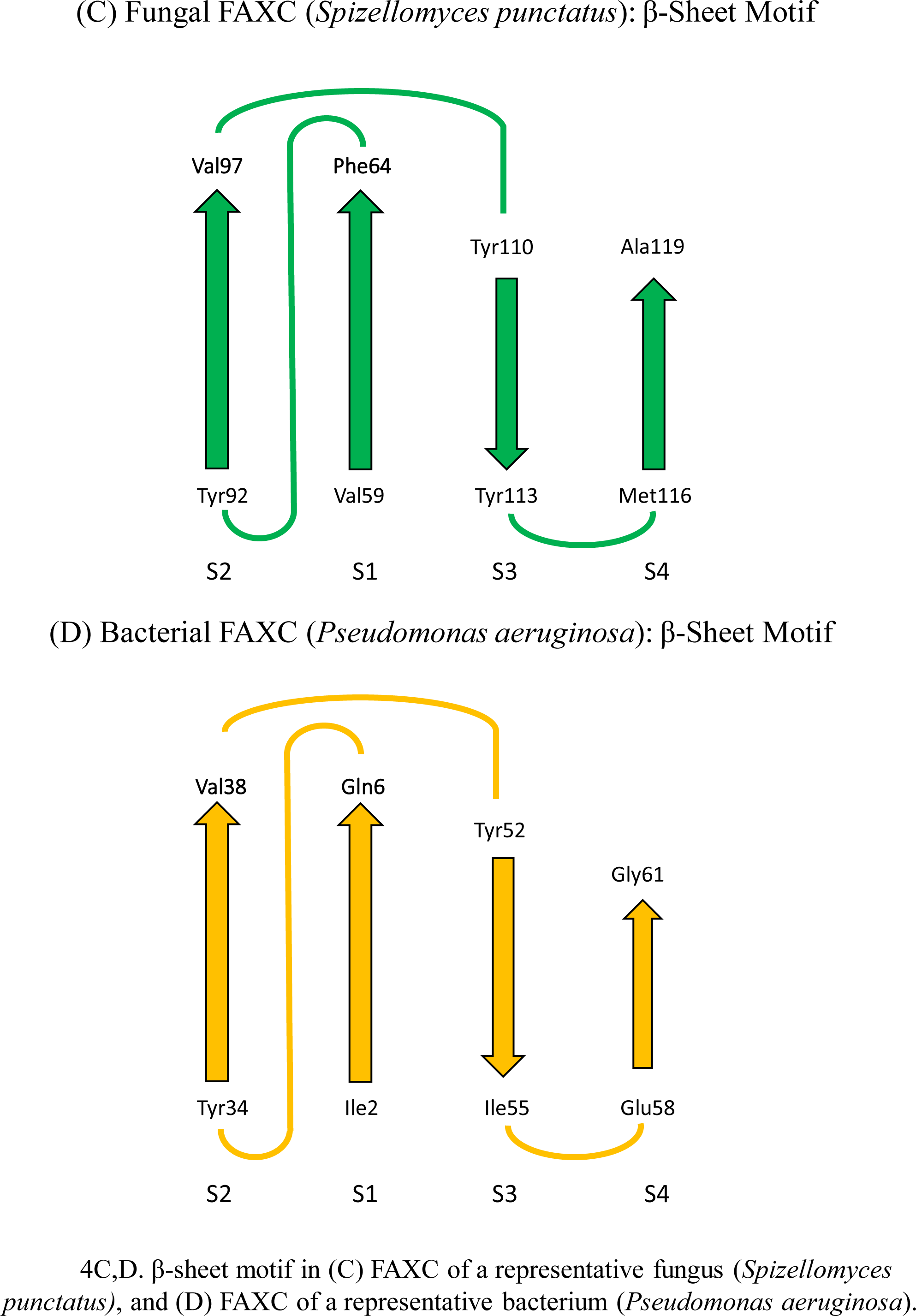
The conserved β-sheet motif of the FAXC proteins of humans, fungi, and bacteria. The motif was detected in the predicted 3D structures of FAXC and metaxin proteins in the AlphaFold protein structure database (Jumper et al., 2021). The figure includes the 4-stranded β-sheet motif in (A) human FAXC isoform 3, (B) human metaxin 3 isoform 3, (C) the FAXC of a representative fungus (*Spizellomyces punctatus*), and (D) the FAXC of a representative bacterium (*Pseudomonas aeruginosa*). The *Spizellomyces* species is a soil fungus that grows in decaying plant matter, while the *Pseudomonas* species is medically significant as a source of hospital-acquired infections. The β-strands form an approximately planar arrangement in the 3D structures, with the order and orientation of the strands S2 (↑), S1 (↑), S3 (↓), S4 (↑). The N-terminal and C-terminal amino acids of each β-strand segment are at the base and tip of each arrow. The 3D β-sheet structure is found in metaxin 1, 2, and 3 proteins, FAXC proteins, and CDR proteins. It is therefore a characteristic feature of metaxin and metaxin-like proteins.

**Figure 5.**
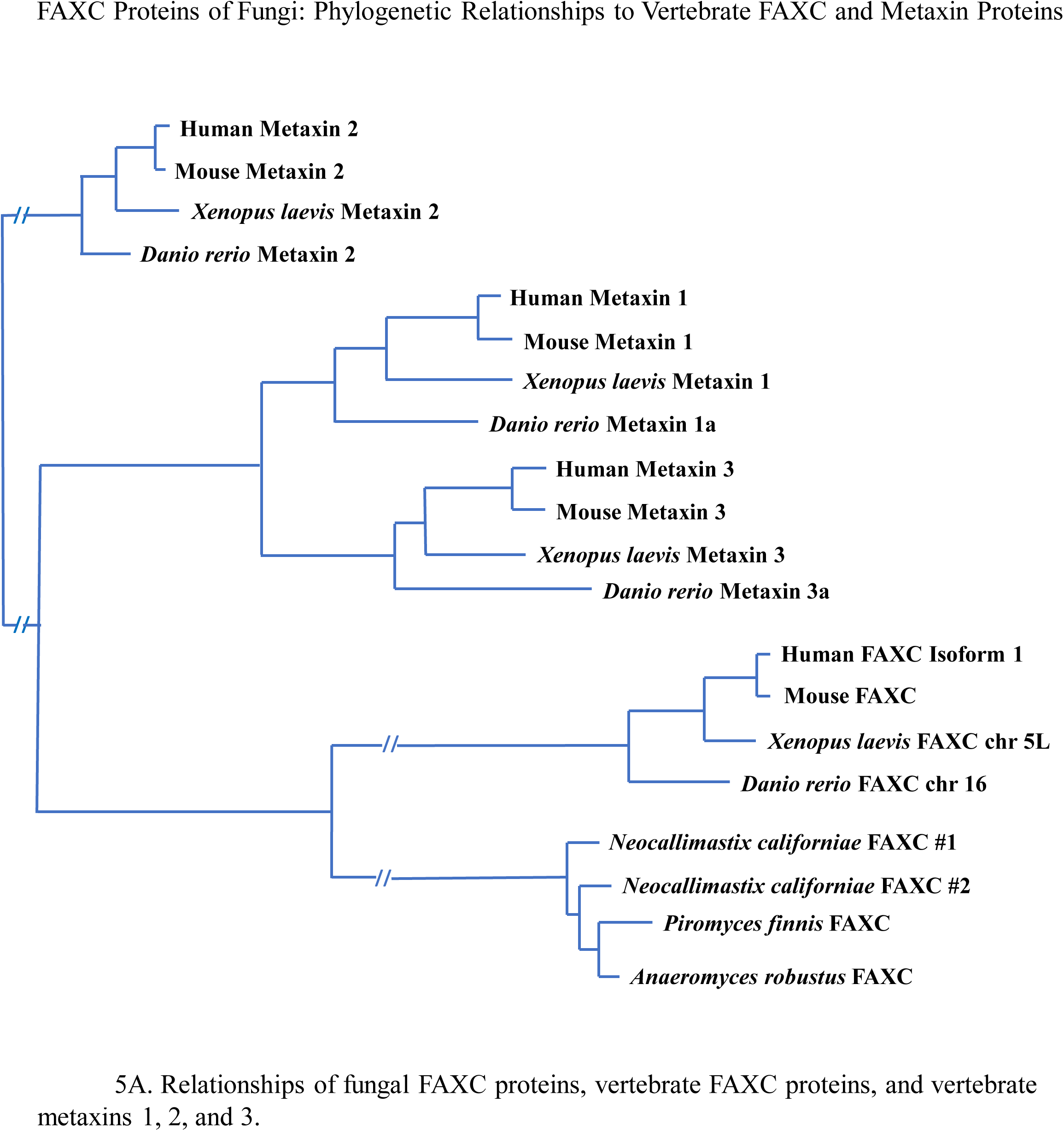
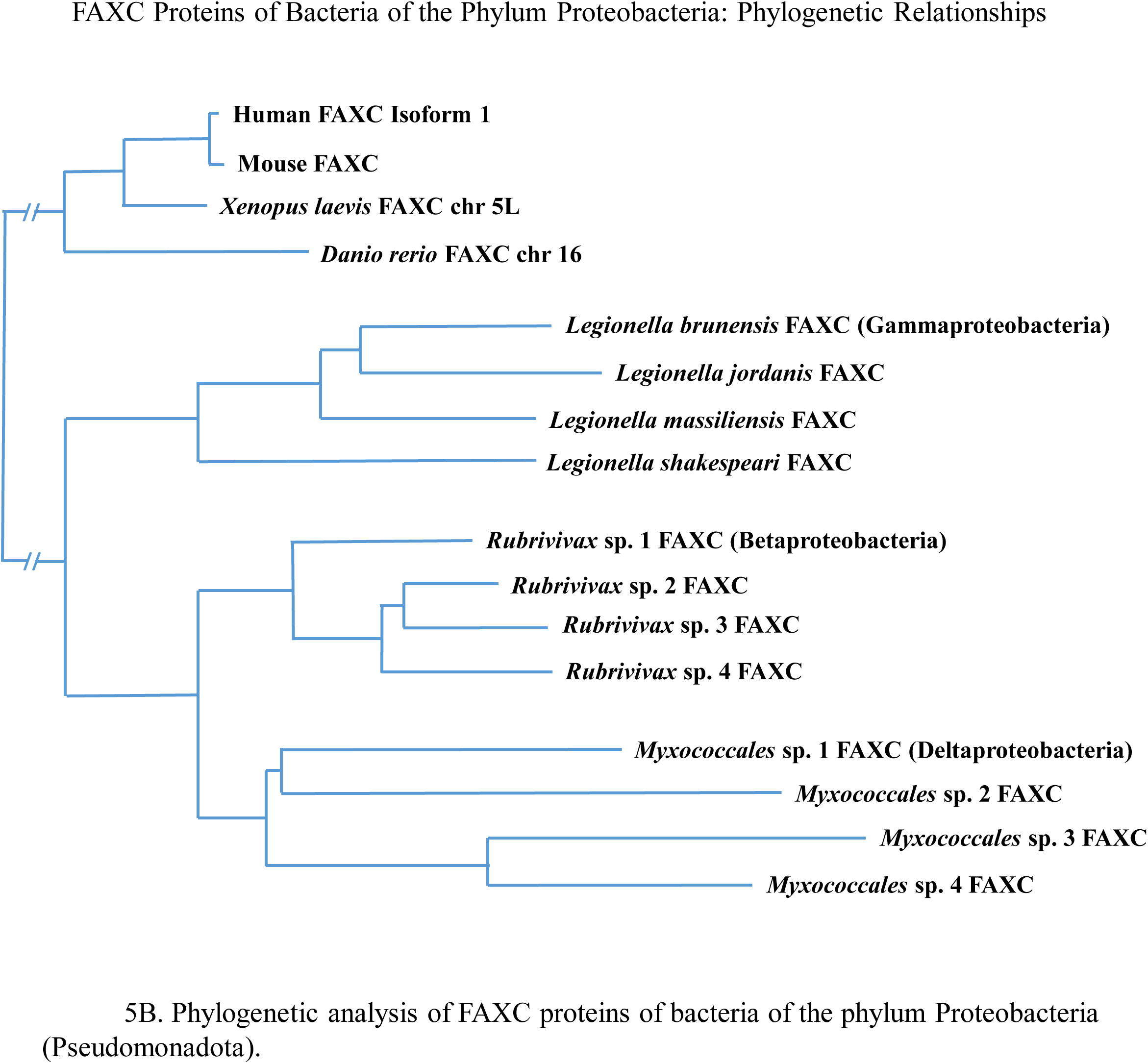
Phylogenetic relationships of fungal and bacterial FAXC proteins with vertebrate metaxin and FAXC proteins. (A) The phylogenetic results in the figure show that fungal and vertebrate FAXC proteins are closely related, while forming two separate groups. The figure also shows that fungal FAXC proteins are related by evolution to vertebrate metaxin proteins. The groups of metaxin 1 and metaxin 3 proteins are closely related to each other. The vertebrate metaxin 2 group is less related to the metaxin 1 and 3 groups as well as the fungal and vertebrate FAXC proteins. These phylogenetic relationships are compatible with the conserved structural features (protein domains, α-helical and β-strand secondary structures) shared by both the FAXC and metaxin proteins. The fungi in the figure are all classified in the Neocallimastigomycota division. (B) The figure shows that bacterial and vertebrate FAXC proteins form separate groups that are related by evolution. The bacteria in the figure represent the Pseudomonadota (Proteobacteria) phylum and the taxonomic classes of the Betaproteobacteria (*Rubrivivax*), Gammaproteobacteria (*Legionella*), and Deltaproteobacteria (*Myxococcales*). The bacterial FAXC groups consist of proteins of bacteria of the same genus but different species, such as the four *Legionella* species. As with the fungal FAXC proteins, the phylogenetic relatedness of the bacterial and vertebrate FAXCs is in keeping with their conserved structural features, as seen in Figures 2, 3, and 4. Bacterial FAXC proteins are also related to vertebrate metaxin proteins, as demonstrated by other phylogenetic results.

**Figure 6.**
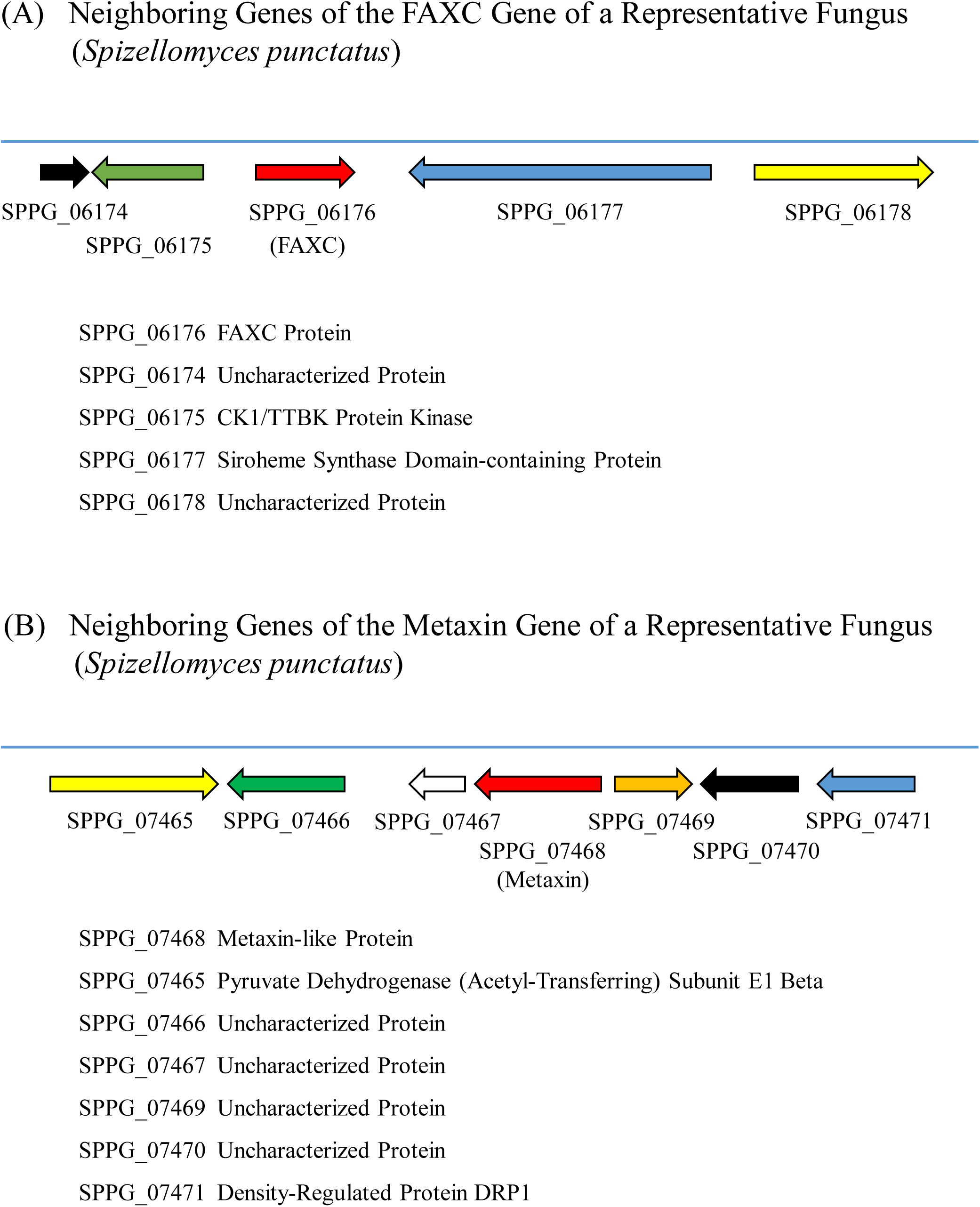
Neighboring genes of the FAXC and metaxin genes of a representative fungus. The representative fungus is *Spizellomyces punctatus*, a soil fungus that gains nutrition from decaying plant matter. Among the genes in Figure 6A that are adjacent to the FAXC gene are the genes for CK1/TTBK protein kinase and siroheme synthase domain-containing protein. The genomic region in Figure 6B for the metaxin gene is different. Genes in this region include those of pyruvate dehydrogenase subunit E1 beta and density-regulated protein DRP1. Genes for uncharacterized proteins are also present.

**Figure 7.**
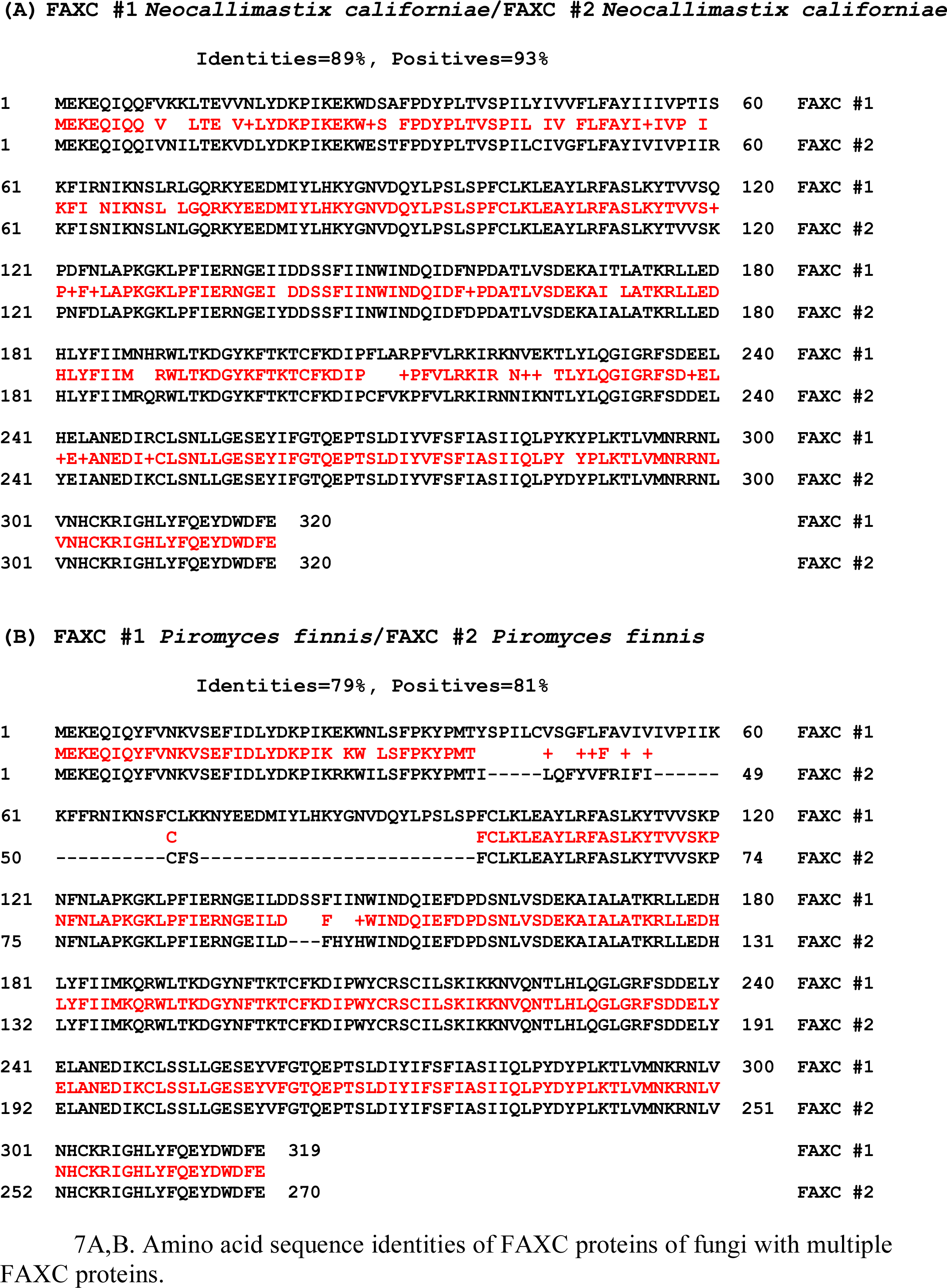
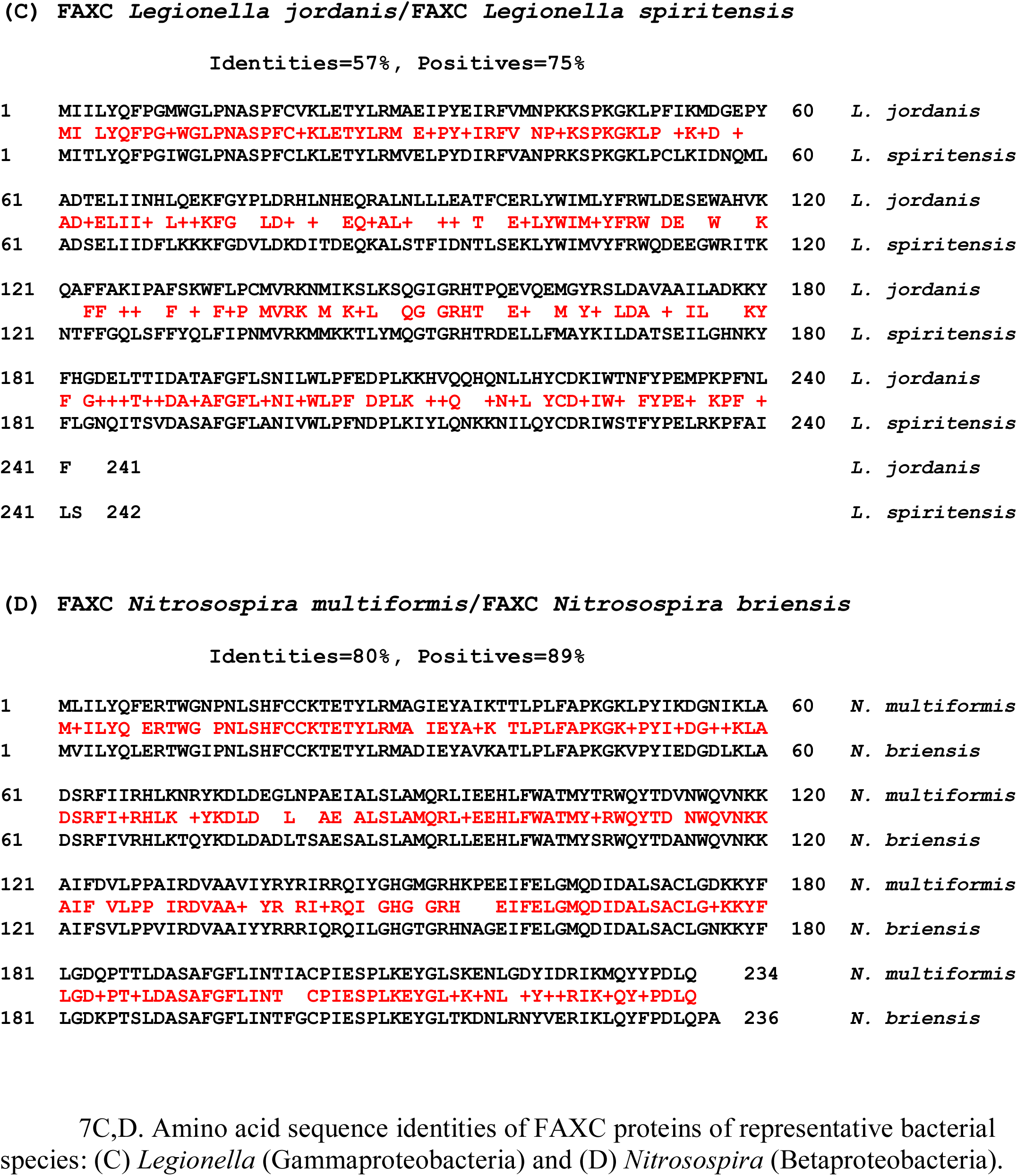
Sequence alignments of multiple FAXC proteins of fungi, and FAXC proteins of different bacteria of the same genus. (A,B) The alignment in (A) includes FAXC #1 and FAXC #2 of *Neocallimastix californiae* (division: Neocallimastigomycota). The identical amino acids are 89% and the positives are 93%. In (B), *Piromyces finnis* FAXC #1 and FAXC #2 are aligned, and show 79% identities and 81% positives. *Piromyces* is also in the Neocallimastigomycota division. The full length sequences of the FAXC proteins in the figure were aligned using NCBI Global Align. The high percentages of identical amino acids indicate that the predicted proteins would have similar secondary and tertiary structures. (C,D) Alignment of amino acid sequences of FAXC proteins of different species of bacteria of the same genus. FAXC proteins of two species of *Legionella* are aligned in (C): *Legionella jordanis* and *Legionella spiritensis*. The identities are 57% and the positives are 75%. The alignment in (D) includes the FAXC proteins of *Nitrosospira multiformis* and *Nitrosospira briensis*. 80% identities and 89% positives are observed. The *Legionella* species are in the Gammaproteobacteria class (Pseudomonadota phylum), while the *Nitrosospira* species are Betaproteobacteria. The high percentages of identical amino acids suggest that the FAXC proteins of different bacterial species of the same genus have similar structural features.

## 3. RESULTS AND DISCUSSION

### 3.1. FAXC and Metaxin Proteins of Fungi: Different Proteins with Similar Structural Features

FAXC proteins and metaxin proteins of fungi are different proteins, as indicated by the low percentages of identities and positives in amino acid alignments (see Figure 1 for examples). But while fungal FAXC and metaxin proteins belong to different categories, other findings, discussed in Sections 3.2, 3.3, and 3.4, show that the proteins have similar conserved structural features. In particular, they have similar protein domains and α-helical and β-strand secondary structures.

In Figure 1A, the alignment shows that the FAXC and metaxin proteins of *Spizellomyces punctatus* share only 17% identical amino acids and 28% positives. *Spizellomyces* is in the Chytridiomycota division. Comparable low percentages, 18% identities and 35% positives, are shown in Figure 1B for the FAXC #1 protein and metaxin protein MTXb of *Heliocybe sulcata* (Basidiomycota division). For FAXC #1 and MTXa of *Heliocybe*, there are just 15% identities and 31% positives. Fungal FAXC proteins frequently occur as multiple proteins, FAXC #1 and FAXC #2. The metaxin proteins of fungi were also found to exist as two proteins, MTXa and MTXb. The soil fungus *Spizellomyces punctatus* is saprotrophic, and derives its nutrition from decaying plants. *Heliocybe sulcata* is a mushroom that is found on decaying wood such as fence posts.

With other fungi, alignments of FAXC and metaxin proteins also demonstrate low percentages of identities and positives, further strengthening the conclusion that FAXC and metaxin proteins are different categories of proteins. Three fungi of the Neocallimastigomycota division are examples. The FAXC #1 and MTXa proteins of *Neocallimastix californiae* have 14% identities and 25% positives, the *Piromyces finnis* FAXC #1 and MTXa proteins have 20% identities and 33% positives, and for *Anaeromyces robustus*, the FAXC and MTX proteins have 20% identities and 35% positives. For two additional fungi of the Chytridiomycota division, similar results were found. *Gonapodya prolifera* FAXC #1 and MTXa have 15% identities and 30% positives, while *Powellomyces hirtus* FAXC #1 and MTXa have 18% identities, 30% positives.

### 3.2. Protein Domains of Fungal and Bacterial FAXC Proteins

A characteristic structural feature of the metaxin proteins is the presence of GST_N_Metaxin and GST_C_Metaxin domains. These domains are found with vertebrate metaxins 1, 2, and 3 and invertebrate metaxins 1 and 2. They are also typical features of the metaxin-like proteins of plants, protists, fungi, and bacteria. Furthermore, the CDR (cadmium-responsive) proteins of *C. elegans* and other nematodes possess the same domains. And, in addition, the FAXC proteins of vertebrates and invertebrates were found to have these domains as prominent structural features (Adolph, 2023).

The FAXC proteins of fungi and bacteria are also characterized by GST_N_Metaxin and GST_C_Metaxin domains. In Figure 2A, the conserved protein domains are shown for examples of fungi of the Neocallimastigomycota, Chytridiomycota, and Basidiomycota divisions. All of the fungi shown have the GST_N_Metaxin and GST_C_Metaxin domains, and almost all have the Tom37 domain. The Tom37 domain was originally described in a yeast protein involved in protein import into mitochondria. The fungal FAXCs frequently exist as multiple proteins, namely FAXC #1 and FAXC #2. For those fungi in Figure 2A that have two FAXC proteins, FAXC #1 is shown.

As Figure 2B demonstrates, the FAXC proteins of bacteria, like the fungal FAXCs, have GST_N_Metaxin and GST_C_Metaxin protein domains as prominent structural features, together with the TOM37 domain. The bacteria in the figure are representatives of major classes of the Pseudomonadota (Proteobacteria) phylum: Alphaproteobacteria, Betaproteobacteria, Gammaproteobacteria, and Deltaproteobacteria. Also included in Figure 2B are the conserved protein domains of human metaxin 1. The similarity of the human metaxin 1 domain structure to the bacterial and fungal FAXC structures is evident. This highlights the metaxin-like nature of the bacterial and fungal FAXCs, as found earlier for vertebrate and invertebrate FAXCs (Adolph, 2023).

Vertebrate FAXC proteins have an additional domain not found with fungal and bacterial FAXCs. This is the FAXC_N domain (Figure 2C). In the figure, FAXC proteins of human, mouse, *Xenopus laevis* (frog), and *Danio rerio* (zebrafish) are all seen to possess the FAXC_N domain. The domain extends from the N-terminal end of the FAXC proteins to the GST_N_Metaxin domain, and has a small overlap with the GST_N_Metaxin domain. The FAXC_N domain has only been found with vertebrate FAXCs and has not been detected with other FAXCs, including those of fungi and bacteria.

### 3.3. Alpha-Helical Secondary Structure of the FAXC Proteins of Fungi and Bacteria

Figure 3A shows the predicted α-helical segments of FAXC proteins of a selection of fungi. The helical segments, which are underlined and in red, are labeled H1 through H8 based on the labeling of human metaxin 1 (HMTX1P). The fungal FAXCs typically have additional helices near the N-terminus of the proteins, while HMTX1P has a ninth helix, H9, near the C-terminus (within [57] in the figure). The fungi represent a variety of fungal divisions: Neocallimastigomycota (*Neocallimastix californiae*, *Piromyces finnis*), Chytridiomycota (*Gonapodya prolifera*, *Powellomyces hirtus*), and Basidiomycota (*Heliocybe sulcata*). The fungal FAXCs shown are all FAXC #1; the existence of two FAXC proteins, FAXC #1 and FAXC #2, is a characteristic of fungal FAXCs.

The pattern of α-helical segments demonstrates that fungal FAXC proteins and human metaxin 1 share an important structural feature – a conserved pattern of α-helices. This supports the idea, along with the conserved β-strand motif (Section 3.4) and conserved protein domains (Section 3.2), that FAXCs are metaxin-like proteins. Like the GST_N_ and GST_C_Metaxin domains in Figure 2, the helices in Figure 3A are defining properties of the FAXC proteins of fungi. The same pattern of α-helices H1 – H8 is found for human metaxin 2 and metaxin 3 in addition to metaxin 1, and is also found for other vertebrate metaxin proteins. Invertebrate genomes encode two metaxin proteins, metaxins 1 and 2, which also display the same pattern of α-helices H1 through H8. A very different group of proteins, the CDR (cadmium-responsive) proteins of *C. elegans* nematodes and other nematodes, also contain α-helical segments H1 – H8. The CDR proteins help to protect against the toxic effects of the heavy metal cadmium. Besides the conserved pattern of α-helices, CDR proteins have the conserved GST_N_Metaxin and GST_C_Metaxin domains found for metaxins and FAXCs.

As Figure 3B demonstrates, bacteria also possess FAXC proteins with a conserved pattern of α-helical segments. As with fungi, the protein domain structures and conserved α-helical segments and β-strand motif (see Section 3.4) of bacterial FAXC proteins are clear evidence that bacterial FAXCs have metaxin-like propeties. All of the bacterial proteins in the figure have the pattern of helices H1 to H8 characteristic of the metaxins, as well as FAXC proteins and CDR proteins. The α-helical segments, indicated in red and underlined, are almost perfectly aligned for the different bacterial FAXC proteins and for human FAXC isoform 1.

Bacterial FAXCs are generally shorter than eukaryotic FAXCs, and don’t have extra helices near the N- and C-termini of the proteins. Four of the bacteria in the figure are in the Pseudomonadota phylum. These bacteria are *Parvibaculum lavamentivorans* (class: Alphaproteobacteria), *Nitrosomonas nitrosa* (Betaproteobacteria), *Pseudomonas aeruginosa* (Gammaproteobacteria), and a Deltaproteobacteria bacterium. The fifth example, *Rhodococcus* sp., is in the Actinomycetota phylum.

### 3.4. β-Sheet Motif in FAXC and Metaxin Proteins of Fungi and Bacteria

Although the predicted secondary structures of the FAXC proteins of fungi and bacteria have α-helical segments as the predominant feature, β-strand segments are also important. They are found as four, short segments labeled S1 – S4 in the N-terminal half of the proteins (Figures 3A and 3B). For fungal FAXCs, there are typically four amino acids in each β-strand segment, with the highest number being six and the lowest number, three. As an example, all four segments S1 – S4 of *Neocallimastix californiae* FAXC #1 in Figure 3A consist of four amino acids. The bacterial β-strand segments are mainly four or five amino acids long. Segments S2, S3, and S4 are in close proximity between helix H1 and helix H2, and segment S1 is before H1 for both fungal and bacterial FAXCs. The β-strand segments of human metaxin 1 (HMTX1P) and human FAXC isoform 1 in Figures 3A and 3B have an arrangement that is similar to that of the fungal and bacterial FAXCs.

The AlphaFold database of predicted 3D structures of proteins (Jumper et al., 2021) revealed that the four β-strand segments are folded to form a three-dimensional β-sheet motif (Figure 4). The structure is found for both fungal FAXC proteins (Figure 4C) and bacterial FAXC proteins (Figure 4D). It is also found in human FAXC protein (Figure 4A) and in human metaxin 3 (Figure 4B). The figure includes human metaxin 3, but the motif is present in human metaxin 1 and metaxin 2, and in the metaxins of other vertebrates and invertebrates. In addition, the four-stranded β-sheet motif is present in the CDR proteins of *C*. *elegans* and other nematodes.

The same 3D arrangement of β-strand segments is observed in all the examples studied. The β-strand segments form a planar structure with individual strands in the order S2, S1, S3, S4. The directionality of the β-strands is ↑ ↑ ↓ ↑. There is some deviation from a strictly planar arrangement particularly for S4 but also for S3. The β-strands are all relatively short, with the greatest lengths commonly being 4 or 5 amino acids. S1 and S2 are the longest, then S3, and finally S4. Variation in the lengths can be seen in the β-strand examples in Figure 4. The N- and C-terminal amino acids of the β-strand segments are indicated at the base and tip of each arrow.

### 3.5. Pylogenetic Relationships of FAXC Proteins of Fungi and Bacteria

Fungal FAXC proteins are closely related to vertebrate FAXC proteins, as revealed by phylogenetic results such as in Figure 5A. Fungal FAXC proteins are also related to vertebrate metaxins, but are not as closely related as fungal and vertebrate FAXCs. The order of relatedness to fungal FAXC proteins is: vertebrate FAXCs > vertebrate metaxin 1 and metaxin 3 proteins > vertebrate metaxin 2 proteins. This is in agreement with the high percentages of amino acid identities (about 45%) in aligning metaxin 1 and metaxin 3 proteins, compared to metaxin 1 (or 3) and metaxin 2 proteins (about 22%). The fungi in the figure (*Neocallimastix*, *Piromyces*, *Anaeromyces*) are in the Neocallimastigomycota division. The vertebrate FAXC and metaxin proteins are from human, mouse, *Xenopus laevis* (frog), and *Danio rerio* (zebrafish). These and additional phylogenetic results along with the shared protein domain structures (Figure 2A and 2C) and conserved α-helical and β-strand segments (Figure 3A) demonstrate the structural relatedness of fungal FAXC and vertebrate metaxin proteins.

For bacteria, Figure 5B shows the evolutionary relationships of FAXC proteins of representative bacteria and vertebrates. Bacterial FAXC proteins are included for three classes of proteobacteria: Betaproteobacteria (*Rubrivivax*), Gammaproteobacteria (*Legionella*), and Deltaproteobacteria (*Myxococcales*). Vertebrate FAXC proteins include FAXCs of human, mouse, *Xenopus*, and zebrafish. The bacterial FAXC proteins form groups of the same genus, and show a close relationship between the different groups. The vertebrate FAXC proteins also show a relationship to the bacterial FAXCs, but a closer relationship exists between the FAXCs of different bacteria.

### 3.6. Adjacent Genes of FAXC and Metaxin Genes of Fungi

The neighboring genes of the FAXC and metaxin genes of a representative fungus are included in Figure 6. The fungus is *Spizellomyces punctatus*, a soil fungus of the Chytridiomycota division that is saprotrophic and obtains nutrition through the extracellular digestion of decayed organic matter. As the figure shows, the genomic regions of the FAXC gene (Figure 6A) and the metaxin gene (Figure 6B) include different genes. The FAXC gene is between the genes for CK1/TTBK protein kinase and siroheme synthase domain-containing protein. In contrast, the genomic region of the metaxin gene contains genes for pyruvate dehydrogenase subunit E1 beta, density-regulated protein DRP1, and the genes for several uncharacterized proteins.

Genome regions with different genes are also found for the FAXC and metaxin genes of humans. The human FAXC gene is next to the *COQ3* (coenzyme Q3 methyltransferase) gene, and other nearby genes include the *USP45* (ubiquitin-specific peptidase 45) gene and the *CCNC* (cyclin 3) gene. In contrast, the genomic region of human metaxin 1 includes the adjacent genes for thrombospondin 3 (*THBS3*) and a glucosylceramidase beta 1 pseudogene (*psGBA1*). Human metaxin 3 is next to the genes for thrombospondin 4 (*THBS4*) and cardiomyopathy-associated 5 protein, while human metaxin 2 has homeobox genes as neighbors.

Other vertebrates have FAXC genomic regions with similarities to the human FAXC region. The zebrafish *Danio rerio*, a widely studied model vertebrate, has two *faxc* genes, *faxca* and *faxcb*, produced by a genome duplication. The *faxcb* gene on chromosome 16 is next to the *coq3* (coenzyme Q3 methyltransferase) gene, and the genomic region also includes genes such as the *usp45* (ubiquitin-specific peptidase 45) gene. However, the *faxca* gene on chromosome 4 has a genomic region with mainly different genes. The frog *Xenopus laevis*, also a model vertebrate, has the *faxc.L* gene on chromosome 5L, with neighboring genes that include the *coq3.L* (coenzyme Q3 methyltransferase L) gene and the *usp45.L* (ubiquitin-specific peptidase 45L) gene.

Invertebrates, for example the model insect *Drosophila melanogaster*, have FAXC genes with adjacent genes that differ from those of both fungi and vertebrates. In particular, the *D. melanogaster fax* gene is between the *Gagr* (GAG-related) and *TMS1* (orthologous to SERINC3, serine incorporator 3) genes. With other invertebrates, the genomic regions of the FAXC genes are generally not conserved and different adjacent genes are the rule. The *C. elegans* FAXC #1 gene, for instance, is next to the gene for trimeric intracellular cation channel type 1B.1, and the genomic region also includes genes for poly(A)-specific ribonuclease and eukaryotic translation initiation factor 4E-5.

### 3.7. Amino Acid Sequence Identities of the Multiple FAXC Proteins of Fungi, and the FAXC Proteins of Different Bacteria of the Same Genus

High percentages of identical amino acids correlate with structural and functional similarities. For fungi, the high degree of amino acid homology for FAXC #1 and FAXC #2 proteins of the same fungal species (discussed below) indicates that the two FAXCs have similar functions. With bacteria, the high percentages of identical amino acids for the FAXC proteins of different species of the same genus (see below) also provide evidence that these FAXCs have similar functions.

Figure 7 includes alignments of pairs of FAXC protein sequences of fungi with multiple FAXCs. In (A), FAXC #1 and FAXC #2 proteins of *Neocallimastix californiae* are aligned and show 89% identical amino acids. In (B), the alignment of FAXC #1 and FAXC #2 of *Piromyces finnis* reveals 79% identical amino acids. Both fungi are anaerobic fungi of the Neocallimastigomycota division. Two FAXCs, FAXC #1 and FAXC #2, have been detected for each. Other examples of fungi that have FAXC #1 and FAXC #2 proteins include fungi of the Chytridiomycota division, *Gonapodya prolifera* and *Powellomyces hirtus*. In addition, *Heliocybe sulcata*, a fungus of the Basidiomycota division, also has genes for FAXC #1 and FAXC #2 proteins. Fungi can also have two metaxin proteins, MTXa and MTXb.

Vertebrates primarily have a single FAXC gene. But there are exceptions. The zebrafish *Danio rerio*, for example, has *faxca* and *faxcb* genes. Invertebrates can have more than 10 FAXC genes. The sea anemone *Exaiptasia diaphana* has at least 13 FAXC genes, and the lancelet *Branchiostoma floridae* has at least 10 (Adolph, 2023).

For bacteria, FAXC proteins of different species but the same genus display high percentages of identical amino acids. This can be observed in Figure 7C for the FAXCs of *Legionella jordanis* and *Legionella spiritensis*, which have 57% identities. *Nitrosospira multiformis* and *Nitrosospira briensis* are included in Figure 7D, and the alignment reveals 80% identical amino acids. *Legionella* is a bacterial genus of the Gammaproteobacteria class of the Pseudomonadota. It includes the major pathogenic species (*Legionella pneumophila*) that causes Legionaires’ disease, a type of pneumonia. *Nitrosospira* is in the Betaproteobacteria class. With other bacteria, a high level of amino acid sequence homology has also been found for the FAXCs of different species but the same genus. For example, amino acid alignments of *Nitrosomonas oligotropha* and *Nitrosomonas ureae* of the Betaproteobacteria show 75% identical amino acids. *Rhodospirillaceae* species #1 and *Rhodospirillaceae* species #2 (Alphaproteobacteria) display 71% identities.

The results presented in this paper provide evidence that the FAXC proteins of fungi and bacteria share structural features with metaxin proteins and can be considered metaxin-like proteins. However, greater experimental evidence is needed to further understand the structure and function of the FAXC proteins of both fungi and bacteria, as well as the FAXCs of vertebrates and invertebrates.

## Notes

### Competing Interest Statement

The authors have declared no competing interest.

